# Adenylyl cyclase 9: fundamental change of regulation in vertebrates and gene sub-functionalization after teleost-specific whole-genome duplication

**DOI:** 10.64898/2025.12.09.693139

**Authors:** Ferenc A. Antoni, Julie Mazzolini, Heather McClafferty, Zhiaho Chen, Laura Szalai, Cristina Xia, Sahad Iqbal, Martin Denvir, András Balla, Dirk Sieger, Michael J. Shipston, Paul Skehel

## Abstract

Adenosine 3’:5’ monophosphate (cAMP) is a ubiquitous signalling molecule generated by the adenylyl cyclase family of proteins which is encoded by ten genes. The biological significance of this diversity is not well understood. In mammals, transmembrane adenylyl cyclase 9 (AC9) is resistant to regulation by heterotrimeric G proteins. A major facet of this resistance is auto-inhibition — in the presence of activated Gsα, AC9 is inhibited by its C-terminal domain. Here, we examined the natural evolution of this seemingly paradoxical control mechanism. At the primary sequence level, the hallmarks of auto-inhibition are apparent in all vertebrates, none are found in invertebrates. Teleost-specific genome duplication (TGD) resulted in *adcy9* ohnologs, one of which lacked the hallmarks of auto-inhibition. This was confirmed in functional assays upon cloning and heterologous expression of the requisite cDNAs. The tissue distributions of the *adcy9* ohnologs in teleost species also pointed to their sub-functionalization. Above all, auto-inhibited *adcy9* was largely restricted to the brain indicating a fundamental role in brain development or function. Our findings document a quantum leap of the regulation of the enzymatic activity of AC9 in vertebrates and the potency of TGD to meet an adaptational challenge through functionally diversified ohnologs.

## Introduction

Signalling through adenosine 3’:5’monophosphate (cAMP) is close to ubiquitous in biological systems. In vertebrates ten genes encode the enzyme of cAMP biosynthesis, adenylyl cyclase (AC). Nine of these give rise to transmembrane proteins (AC1 to AC9) while one gene product is a soluble enzyme (Dessauer et al. 2017; Antoni 2020; Ostrom et al. 2022). All transmembrane ACs (tmAC) conform to the same blueprint of a single polypeptide chain of quasi-symmetrical design, in which two sets of six closely-spaced transmembrane domains are preceded, linked, and terminated by long cytoplasmic and relatively short extracellular loops (Fig. 1a). High-resolution cryo-electronmicroscopic maps of AC5, AC8 and AC9 all show that tmACs consist of three structurally distinct, functionally connected domains (Fig 1b) (Schuster et al. 2024). The transmembrane arrays are closely packed and interfaced in the cellular lipid bilayer. Transmembrane helices 6 and 12 face each other and continue into the cytoplasm as alpha-helices (signalling helix 6 and 12, respectively) forming a short parallel coiled-coil before the start of the C1a_C2a catalytic core. The isoform specific N cytoplasmic loop, the C1b and most of the C2b domain could not be resolved by cryoEM, most plausibly because these are intrinsically disordered domains (Suppl fig 1.). Importantly, a short segment of the C2b domain could be detected yielding the striking finding that in the presence of activated Gsα, the active site of the enzyme is occupied by a previously identified short autoregulatory motif (ARM) (Pálvölgyi et al. 2018) comprised of 15 amino acid residues (Qi et al. 2019; Nomura et al. 2025). Removal of the ARM markedly enhanced the stimulation of AC9 by Gsα in a cell free system (Qi et al. 2019) as well as its response to the activation of receptors coupled to Gs in intact cells (Pálvölgyi et al. 2018; Chen and Antoni 2023).

**Figure 1.**
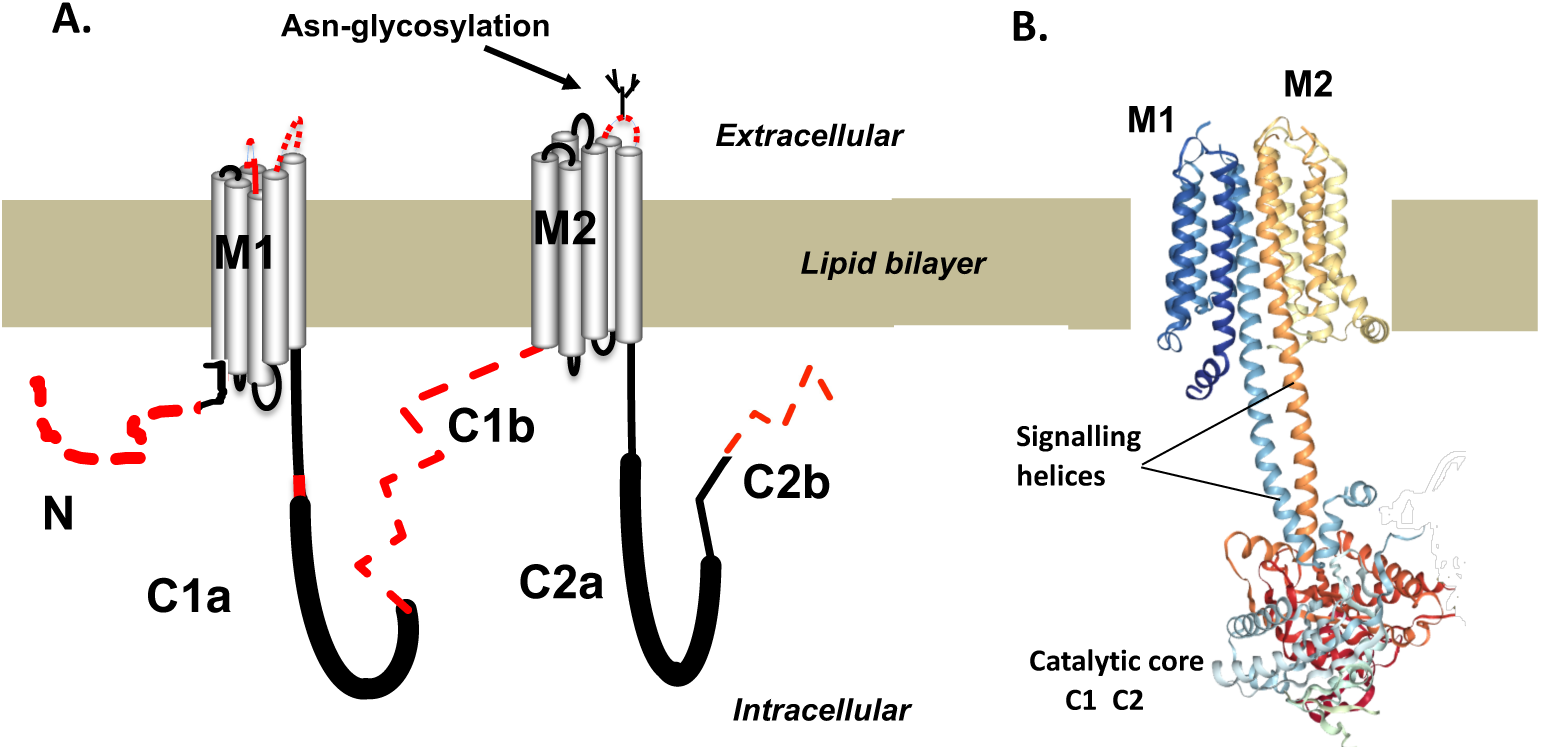
Schematic representations of adenylyl cyclase 9 (AC9). A) A single polypeptide with two six-member transmembrane (M1 and M2) arrays. Dashed red lines indicate where the cryoEM failed to clarify the structure. Thick lines indicate the catalytic core that is similar in all paralogues of AC9. N – amino-terminal domain, C1b and C2b are isoform-specific cytoplasmic loops. B) The 3D structure of AC9 obtained by cryoEM. Note there are three structurally distinct regions: the transmembrane arrays, the signalling helices and the cytoplasmic catalytic domain formed by C1 and C2.

An ancestral AC9 (*adcy-1* gene) is already present in *C. elegans* (Korswagen et al. 1997; Berger et al. 1998). This enzyme has virtually no C2b domain and is readily stimulated by Gs coupled receptors as well as forskolin (Berger et al. 1998; Mano and Driscoll 2009; Saifee et al. 2011). Thus, the marked autoinhibition of physiological stimulation in mammalian AC9 orthologues appears to be a veritable biological paradox worthy of further analysis with particular reference to its origin during the course of natural evolution.

## Materials and Methods Database searches

The databases and software packages provided by NCBI (https://www.ncbi.nlm.nih.gov), European Bioinformatics Institute (https://www.ebi.ac.uk), ENSEMBL (https://www.ensembl.org/index.html), PhyloFish (http://phylofish.sigenae.org/index.html) and Zebrafish information network (https://zfin.org/) were used to collate genomic data and carry out sequence alignments. With respect to sequence alignments, all of the sequences used were designated as adenylyl cyclase 9-like by the database curators on the basis of BLAST max scores and had error rates (E) of 0≤E<10^-50^. Protein sequence alignment and phylogenetic tree construction was carried out with COBALT (https://www.ncbi.nlm.nih.gov/tools/cobalt/cobalt.cgi).

### Zebrafish care and husbandry, ethical approval

All experiments were carried out in accordance with the accepted standards of humane animal care under the regulation of the UK Animal (Scientific Procedures) Act 1986 and EU Directive 2010/63/EU; all experiments were approved by the University of Edinburgh Animal Welfare and Ethical Review Board.

Adult wild-type fish were housed according to standard operating procedures (https://zfin.org/zf_info/zfbook/zfbk.html, (Brand et al. 2002)) and maintained with a 14h light:10h darkness photoperiod cycle at an ambient water temperature of 28.5°C. The established line used was WIK. Adult zebrafish were housed in 10L tanks at a density of 2–3 fish per litre (maximum of 30 fish per tank).

All larvae used in these experimental studies were under the age of 5 days post fertilisation; embryos were collected from random matings and then correctly staged for development, according to (Kimmel et al. 1995).The larvae were kept at 28 °C on a 14 hours light/10 hours dark photoperiod. Embryos were obtained by natural spawning from adult *tg(mpeg1:GFP*) (Ellett et al. 2011) and wild-type (WIK) zebrafish strains. Embryos were raised at 28.5°C in embryo medium (E3) and treated with 200 μM 1-phenyl 2-thiourea (PTU) (Sigma) from the end of the first day of development for the duration of the experiment to prevent pigmentation.

### Macrophage isolation and FACS analysis

Macrophages were isolated by FACS from heads of mpeg1:GFP 5 days post fertilization (d.p.f.) larvae as previously described (Mazzolini et al. 2018). FACS allowed cell separation from debris in function of their size (FSC-A) and granularity (SSC-A). Single cells were then separated from doublets or cell agglomerates (FSC Singlet; SSC Singlet). From the single-cell population, a gate was drawn to separate live cells (DAPI−) from dead cells (DAPI+). Unstained cells from wild-type (WIK) zebrafish strain were used as controls to draw gates corresponding to macrophage (GFP^+^) populations segregated from the live cell population gates. FACS data were analyzed using FlowJo Software (Treestar, Ashland, OR).

### RNA extraction and cDNA amplification

Total RNA extraction from FACS derived macrophages was performed using the Qiagen RNeasy Plus Micro kit according to the manufacturer’s guidance (Qiagen). For qPCR, RNA sample quality and concentration were assessed using the LabChip GX Touch Nucleic Acid Analyzer and RNA Pico Sensitivity Assay. All RNA samples with a RIN score > 7 were transcribed from the same amount of RNA into cDNA using the Super-Script® III First-Strand Synthesis System (Invitrogen).

### Quantitative PCR for *adcy9* expression in zebrafish

Tissues from adult or 5 d.p.f. zebrafish (Wik) were excised after an overdose of MS-222 (tricaine methanesulfonate - 40 μg ml^−1^, Sigma), and placed in “RNA later” (Ambion, Life technologies) solution. The RNA was extracted with ReliaPrep™ RNA Miniprep kit (Promega) as per the manufacturer’s protocol and reverse transcribed using SuperScript™ IV First-Strand Synthesis System (ThermoFisher). Approximately 100-700ng RNA was primed with a 1:1 mix of oligo-dT and random hexamers and relative gene expression was determined by comparative qPCR using Power SYBR™ Green PCR Master Mix (Applied Biosystems) on a StepOnePlus™ Real-Time PCR System. Primers for the zebrafish *adcy9* coding sequences were designed by Oligo Tech Service (Sigma/ DORSET /UK). The *adcy9* genes of zebrafish are found on chromosome 3 and 22.

Chr3 sense: TGTGAGTTCAGGACTTAC Chr3 antisense: TTAGATGTACCATTGCTTAC 95.9% efficiency on zebrafish brain over 3 dilution factors. Chr22 sense: TACACAACCTTCCTCAAT, Chr22 antisense: TCTGCTTCATCTTCTTCT 94.2% efficient on total zebrafish cDNA over 3 dilution factors. The house-keeping gene *bactin2* was used as the reference (Casadei et al. 2011).

### Partial characterization of zebrafish AC9 isoforms

The full length cDNAs of the zebrafish AC9s could be amplified from cDNA prepared from whole 5 d.p.f. larvae with Phusion polymerase (NEB, Hitchin,UK) using primers

> Chr3Fwd 5’ATGGCTTCTCCTCAGCATCTAC3’ Chr3Rev 5’TCAAAGTTTTGTCATTTCACCCA3’ Chr22Fwd 5’ATGGCATTGCCAGAGCAACAGC3’ Chr22Rev 5’TCACTCCACTGCCCTGAGCTTTG3’

These primers were subsequently extended by the requisite 15 bases and used to amplify the full length AC9 by Phusion PCR for insertion into pEGFP_C2 (TakaraBio) by Gibson assembly (Gibson et al. 2009) as implemented in the TakaraBio InFusion cloning kit. The carboxyl-terminal C2b domains of both AC9s were removed by PCR with the Phusion polymerase/Infusion procedure resulting in AC9a_C2a and AC9b_C2a. All AC9 cDNA plasmids were verified by restriction digests and Big Dye or nanopore DNA sequencing.

Human embryonic kidney 293 FT (HEK293) cells were maintained and propagated as previously reported (Antoni et al. 1998; Pálvölgyi et al. 2018). Intracellular levels of cAMP were monitored by light emission from the cAMP biosensor protein Glosensor22F (Binkowski et al. 2011) as described previously (Chen and Antoni 2023). Briefly, the cells were transfected in suspension (10^6^ cells /ml) with up to 1.33 µg plasmid of interest plus 0.67 µg of Glosensor 22F DNA, with 1.6 µl of Lipofectamine 2000 (Life Technologies) reagent per ml of cell suspension in 6cm diameter culture dishes (Greiner). Forty-eight h after the start of the transfection the cells were replated at 10^5^ cells/well in poly-L-lysine coated 96-well white tissue-culture plates (Greiner). After 24h the culture medium was changed to a serum free mixture of (1:1) Dulbecco’s MEM and Ham’s F12 medium and the cells were incubated in 5% CO2 atmosphere at 37°C for 1h. Subsequently, the medium was changed to Hank’s balanced salt solution containing 1 mM MgCl_2_, 1.5 mM CaCl_2_, 10 mM HEPES pH7.4, 1mM luciferin and the plates were transferred to a plate warmer set to 32°C. After 60 min the recording of bioluminescence was started in a Lumistar Omega (BMG Labtech) plate reader at 32°C. Basal levels were recorded for 30 min after which isoproterenol was added by a hand-held multichannel pipette. The response to isoproterenol was monitored for 30 min after which forskolin 5µM/IBMX 5mM was added and the recording terminated 30 min afterwards. The cAMP levels elicited by the forskolin 5µM/IBMX 5mM mixture plateaued after 5 min and the average plateau relative light unit (RLU) value was used to standardize the cAMP production in each well to obviate the inherent variability of transient transfections. This standardization is possible because AC9 is not stimulated by 5 **μ**M forskolin even when stimulated by Gs**α,** and there is a strong synergy between Iso and forskolin to stimulate the HEK293 cells’ endogenous adenylyl cyclases (Chen and Antoni 2023). The standardized RLU data were analysed after log-transformation by one-way ANOVA followed by Tukey’s multiple comparisons test as implemented in Graphpad Prism, v.6.0.

### Immunoblots and confocal microscopy

HEK 293 cells transiently expressing the zebrafish eGFP-AC9 conjugates were processed for immunoblots as previously reported (Pálvölgyi et al. 2018) except that whole cell extracts were analysed. The blots were reacted with rabbit antibodies against GFP (TakaraBio), followed by anti-rabbit IgG Fab fragments conjugated to Alexa 685 dye (Invitrogen, UK), and subsequently scanned in a LiCor Odyssey fluorescence imager. For confocal microscopy the transiently transfected cells were processed as previously described (Pálvölgyi et al. 2018) and imaged in a Zeiss confocal microscope detecting the fluorescence signal from eGFP.

## RESULTS

### ARM is specific to the vertebrate subphylum

A *blastp* search with residues 1243-1353 of human AC9 (the ARM motif is found at residues 1261-1273) returned 1579 hits. Alignment of these sequences showed a close to 100% conservation of the ARM (Suppl Fig 2). The main consistent change found was at position 1264 (to A or I), and was restricted to ray-fin fish. Upon exclusion of vertebrates no hits other than synthetic constructs were found either using the DNA sequence with the *megablast* protocol or the protein sequence with settings of *tblastn* to find less similar sequences.

The phylogenetic tree of selected AC9 protein sequences is shown in Suppl Fig. 3. The node of the split between invertebrates and vertebrates is clearly apparent.

A comparison of human AC9 with the AC9 orthologues of *tunicata,* the invertebrate subphylum most closely related to vertebrates, is shown in Fig 2. The most conspicuous change is the appearance of the long C2b domain harbouring the ARM. Moreover, modifications at positions N515, L519, Q522, L523 (human AC9 numbering) of 4-alpha helix of the C1a domain, the main element of the forskolin binding-pocket are evident. Further relevant changes are at Y1082 and A1112, both previously shown to influence the response of mouse AC9 to forskolin (Yan et al. 1998). The changes listed above are 100% conserved in the AC9 orthologues of vertebrates. The AC9 orthologues of cyclostomi such as the hagfish (*Eptatretus burger)*, the sea-lamprey (*Petromyzon marinu)* and the arctic lamprey (*Lethenteron camtschaticum* also known as *Lethenteron japonicum)*) already feature the full complement of the changes listed above for human AC9 (Suppl Fig 4). Specifically, the long C2b domain is present and includes the highly conserved ARM and its immediate sequence environment. The contact sites for ARM (Qi et al. 2019) in 4α-helix of the C1a catalytic domain have the four AC9 specific residues. In addition, the low forskolin response configuration (Tang and Hurley 1998; Yan et al. 1998) at Y1082 and A1112 in human AC9 is also conserved in all three of the cyclostome orthologues. Taken together, the hallmarks of autoregulation in the primary structure of AC9 are already apparent in the oldest species of currently living vertebrates. Moreover, the ARM as a highly conserved segment of a long, isoform-specific C2b domain of AC9 appears to be specific to and highly conserved in the vertebrate subphylum.

**Figure 2.**
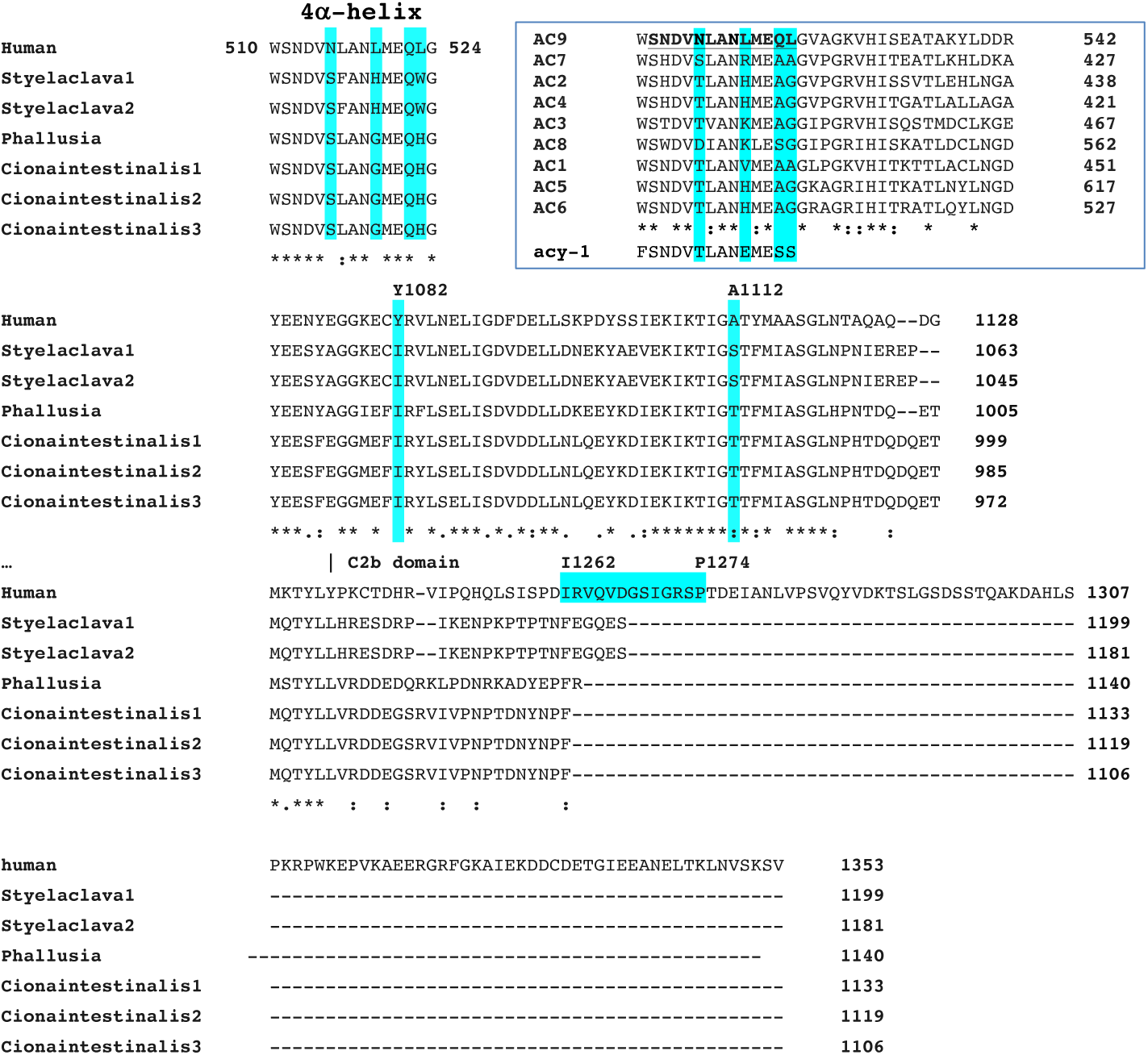
Alignment of relevant sections of the primary sequence of human adenylyl cyclase 9 with its predicted orthologues from *tunicata,* the invertebrate species phylogenetically closest to vertebrates (see also Suppl Fig 3 for justification) . Numbering refers to human AC9. Note: 1) The lack of a sizeable C2b domain in *tunicata; 2)* The presence of three residues in the 4-α helix that are unique to the human enzyme. In fact, these are 100% conserved vertebrate orthologues of AC9; 3) The forskolin responsive configuration in *tunicata* at the homologs of human AC9 Y1082 and A1112. **INSERT:** Shows the alignment of human AC paralogues in the area of the 4-α helix (in bold) in the C1a domain. Note residues unique to AC9 highlighted in cyan, which are fully conserved in the orthologues of AC9 of vertebrate species. At the bottom, acy-1 is the homologous segment of the AC9 orthologue of *C. elegans*, which is responsive to forskolin. It has closer sequence similarity with the non-AC9 residues than with AC9.

### Teleost fish have *adcy9* ohnologs

Searches with *tblastn* for the entire human AC9 primary sequence revealed that several ray-fin species have two AC9 genes. Ray-fin species such as the bowfin (*Amia calva*) and the spotted gar (*Lepisosteus oculatus*) that appeared prior to teleost specific genome duplication (TGD), (see (Braasch and Postlethwait 2012; Pasquier et al. 2017) for reviews) have a single AC9 gene (*adcy9a*) encoding a protein with an ARM that closely resembles the human version (Suppl. Fig. 5). Currently living teIeosts have undergone TGD (e.g. the zebrafish) and have at least two *adcy9* genes. That these are ohnologue pairs has been established for at least four species including zebrafish and medaka (Singh and Isambert 2020). There is invariably an *adcy9a* gene that contains the ARM, and usually has some subtle modifications most notably V1264A within ARM and W1158G in the C2a catalytic domain, when compared to the mammalian and pre-TGD counterparts of fish. The latter mutation affects a W residue unique to AC9 that participates in the binding of activated Gsα to bovine AC9 (Qi et al. 2022). Tryptophan at this position is shared by all mammalian AC9 orthologues. More significantly, the ohnologue *adcy9b* encodes a cyclase protein (AC9b) that has a long C2b domain, but lacks the ARM sequence. In AC9b, the residues specific to mammalian AC9 listed in the previous section are in the configuration found in AC9a – i.e. low forskolin response*. Salmo trutta*, and goldfish (*Carassus auratus*) which have undergone salmonid and cyprid specific genome duplications after TGD, respectively (Postlethwait 2007), have four *adcy9* genes on different chromosomes, two each of *adcy9a-*like and *adcy9b*-like genes, respectively.

A different set of *adcy9* genes is found in the sterlet (*Acipenser ruthenus*). The two *adcy9* genes on chromosomes 13 and 22, respectively, contain the ARM i.e. are *adcy9a*-like. Similarly, in paddlefish (*Polydon spatula*), there are two *adcy9a*-like genes on chromosome 18 and 26 (suppl. Fig. 6). The post-TGD modifications of V1264A within ARM and W1158G in the C2a catalytic domain are not present in any of the predicted proteins (suppl. Fig. 6). These findings are consonant with the report showing the ancestor of the sturgeon-paddlefish lineages had undergone whole genome duplication and indicate that the *adcy9* genes have remained unaltered after the separation of the sturgeon and paddlefish lineages (Redmond et al. 2023). Taken together, TGD has resulted in the ohnologue genes *adcy9a* and *adcy9b.* The primary sequences of the encoded proteins are distinguished by the presence or absence of the ARM in the long C2b domain. The phylogenetic tree (Suppl Fig. 3) of AC9 highlights that there are at least two lineages in teleosts. These can be readily discerned by the C2b domain of AC9b: BLAST searches with the respective C2b domain sequences of AC9b of *D. rerio* and *O. latipes* gave different, non-overlapping sets of hits.

### Differential expression of *adcy9a* and *adcy9b* in tissues indicate sub-functionalization

The Phylofish database “Depth View” feature was used to assemble a semi-quantitative comparison of the distribution of the two *adcy9* gene-transcripts in the tissues of post-TGD teleosts on the basis of the number of mRNAseq hits. Importantly, *adcy9a* mRNA was largely restricted to the brain while *adcy9b* mRNA predominated in all peripheral tissues examined (Fig 3A and B). Follow-up analysis by RT-qPCR of selected zebrafish tissues supported this notion: of the tissues examined only the brain – adult as well as 5 d.p.f. larva — showed relatively high levels of *adcy9a* mRNA (Fig 3C). Figure 3D shows a time-line and the actual mRNAseq counts for the spotted gar, zebrafish and medaka. Note the relatively high levels of expression of *adcy9a* in the brain for the teleost species, and the very low levels in the heart.

**Figure 3.**
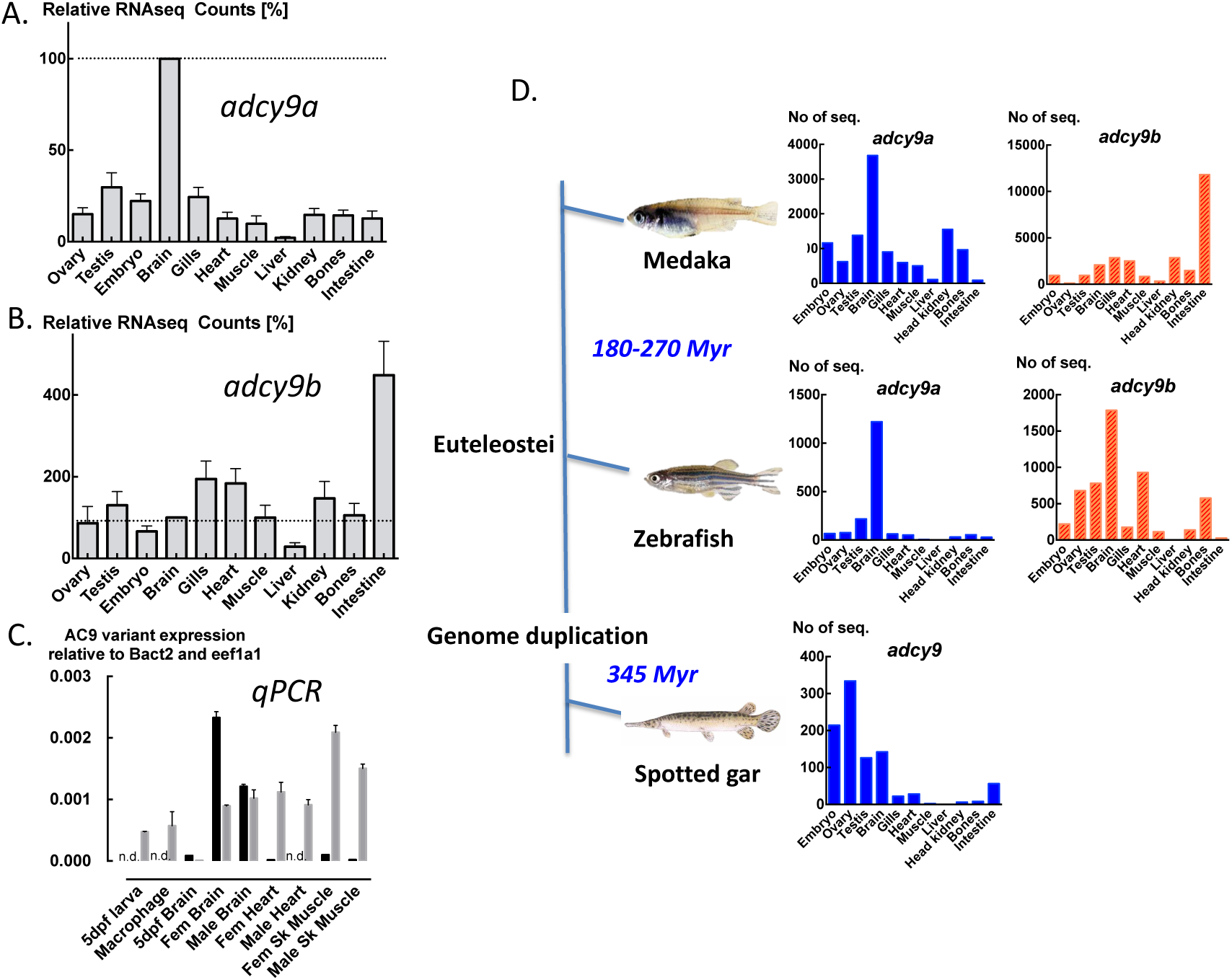
Expression of the *adcy9* genes in the tissues of bony fish. A and B: RNA seq hits from the PhyloFish database. The values are expressed as a percentage of the number of respective hits from brain. For *adcy9a* this was 2623±355, for *adcy9b* 845±184 Mean±S.E.M. N=14. C: RT-qPCR data for the relative abundance of *adcy9a* and *b* mRNAs from zebrafish tissues and 5 d.p.f. whole embryos – filled columns *adcy9a*, empty columns *adcy9b*. Data are Means±S.D., n=5 technical replicates per group. D: The time-line of fish through TGD. The graphs represent the numbers of RNA seq hits from the PhyloFish database for *adcy9a* and *b*.

### Distinct features of cAMP production by the AC9 paralogues of zebrafish

In the case of the zebrafish, *adcy9a* is on chromosome 3 and *adcy9b* is on chromosome 22. The zebrafish *adcy9a* has not been unambiguously identified by exon prediction methods (NCBI or Ensembl servers). However, a full-length transcript (FDR_LOC101477119.1.1) is found in the PhyloFish dataset (http://phylofish.sigenae.org/index.html) and all ten exons could be identified in the chromosome 3 assembly (reverse strand, position 9739058 to 12113782) of the Ensembl server. The cDNAs of both zebrafish AC9 isoforms were cloned from first strand cDNA prepared from 5 d.p.f. embryos. The aligned primary sequences are shown in Fig 4. It is clear that these isoforms are similar, sharing the “low forskolin response” configuration of vertebrate AC9 orthologues. The catalytic domains are near identical. The post-TGD specific modification W1158G is present in both paralogues and AC9a also features the common post-TGD modification V1264A of the ARM. As for other AC9b orthologues, a double leucine motif in the N terminal domain L9L10, that is reportedly responsible for receptor-induced internalization of mammalian AC9 (Lazar et al. 2020), is absent from zebrafish AC9b, but present in AC9a.

**Figure 4.**
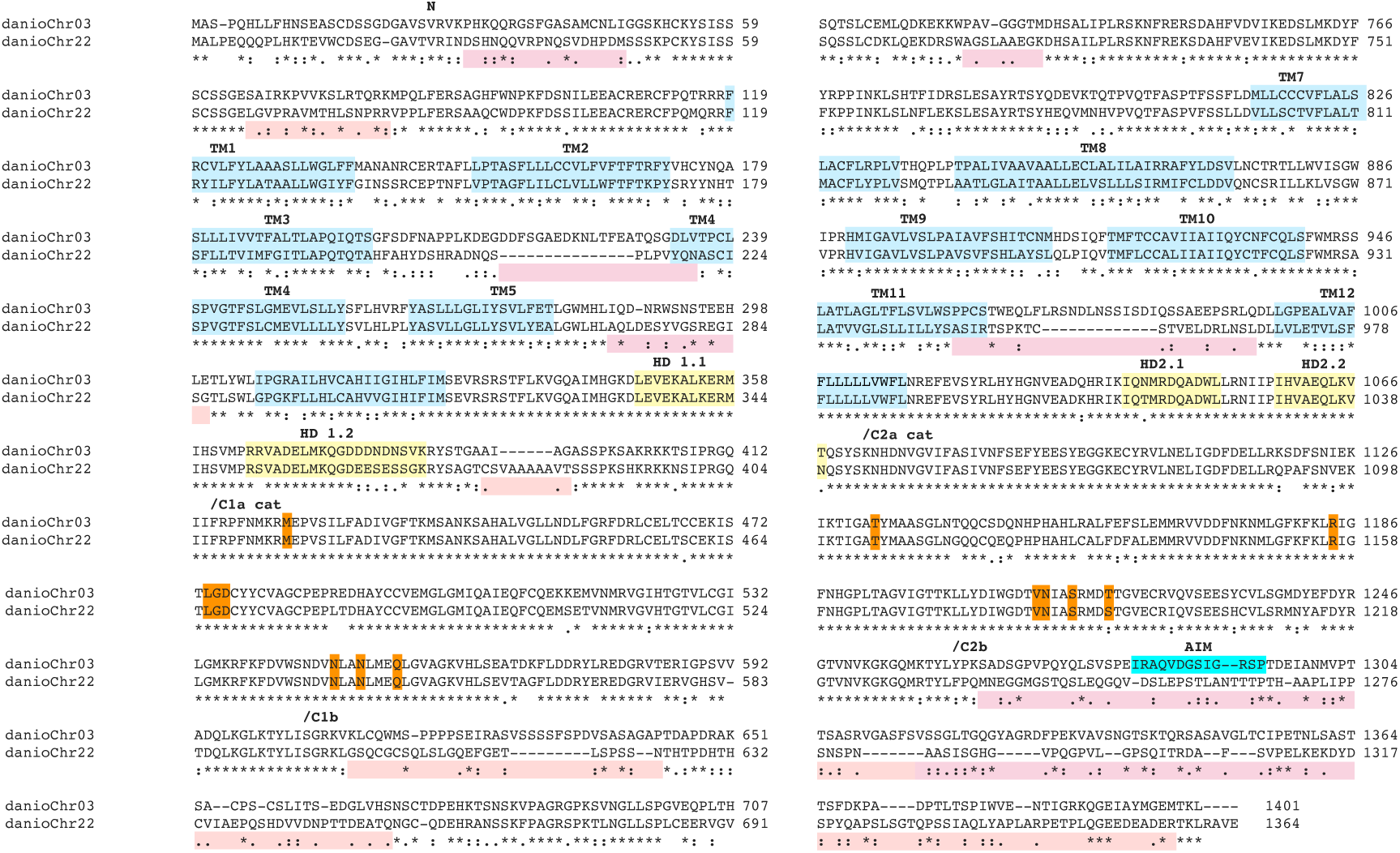
Alignment of the primary structures of the AC9 paralogs of zebrafish. N – N terminal cytoplasmic domain, TM - transmembrane helices (light blue), HD1.1 , 1.2 , 2.1, 2.2 - alpha-helical domains in the signalling helix (yellow). Areas of significant sequence variation are highlighted in pink. The residues in C1a and C2a highlighted with orange interact with the ARM (highlighted in cyan) when it occludes the active site of the bovine AC9.

Both AC9 cDNAs were transiently expressed with eGFP fused in frame to the N-terminus in HEK293FT cells. Western blots of membranes prepared from transiently transfected HEK293FT cells showed similar levels of expression for all AC9 proteins (Fig 5A). Visualisation of eGFP fluorescence in HEK 293 cells transfected with zebrafish eGFP-AC9 fusion proteins indicated that the majority of fluorescent cells expressing zebrafish eGFP-AC9a showed labelling at the perimeter of the cells (Fig 5B, left panel). By contrast, eGFP-AC9b was also present at significant levels in the cytoplasm including perinuclear densities of immunoreactivity (Fig 5B, right panel).

**Figure 5.**
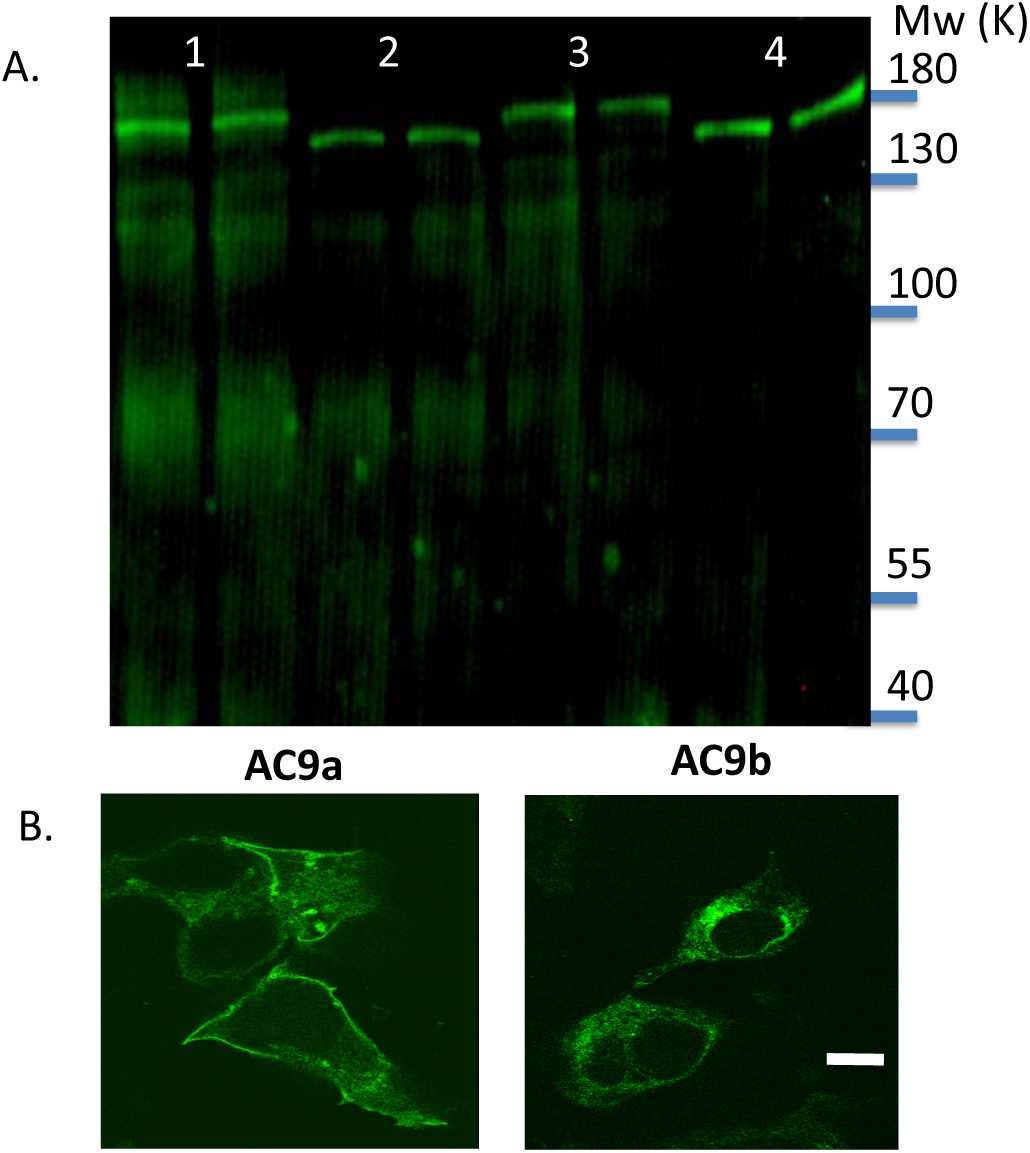
Heterologous expression of the zebrafish GFP-AC9 paralogs in HEK293FT cells. A. Western blot of crude membrane extracts reacted with anti-GFP serum. Samples (in duplicate) are: eGFP-AC9a (1) eGFP-AC9a_C2a (2) eGFP-AC9b (3) eGFP-AC9b_C2a (4) Note lower apparent molecular weight of the C2b truncated versions AC9a_C2a and AC9b_C2a. B. Distribution of eGFP fluorescence in HEK293 cells transiently transfected with plasmids encoding eGFP-AC9a or eGFP-AC9b. A 1µm thick confocal image slice is shown. The scale bar represents 10µm.

Intracellular cAMP levels were monitored by the detection of light emitted by co-transfected glosensor22F. Comparison of AC9a and AC9b showed that the former produced significantly higher basal levels of cAMP-dependent light emission (Fig 6A). Activation of β2-adrenergic receptors endogenously expressed by HEK293FT cells (Cullum et al. 2023) by isoproterenol (Iso), produced concentration dependent increases of cAMP. Notably, the Iso-induced cAMP response produced by AC9a was not significantly different from that of the groups expressing the backbone vector pcDNA3.1 (Fig 6B). This is similar to previous findings with human AC9 (Pálvölgyi et al. 2018; Chen and Antoni 2023). In contrast, a robust rise of intracellular cAMP in response to isoproterenol which was well above the host cell response occurred in the cells expressing AC9b (Fig. 6B-D). These findings strongly indicated autoinhibition by the C2b domain through ARM in AC9a as previously observed for human and bovine AC9 (Pálvölgyi et al. 2018; Qi et al. 2019; Chen and Antoni 2023). Accordingly, when the C2b domain was removed from AC9a, the response to Iso was not different from that observed with AC9b. In contrast, removal of the C2b domain slightly diminished the Iso response of AC9b (Fig 6B-D).

**Figure 6.**
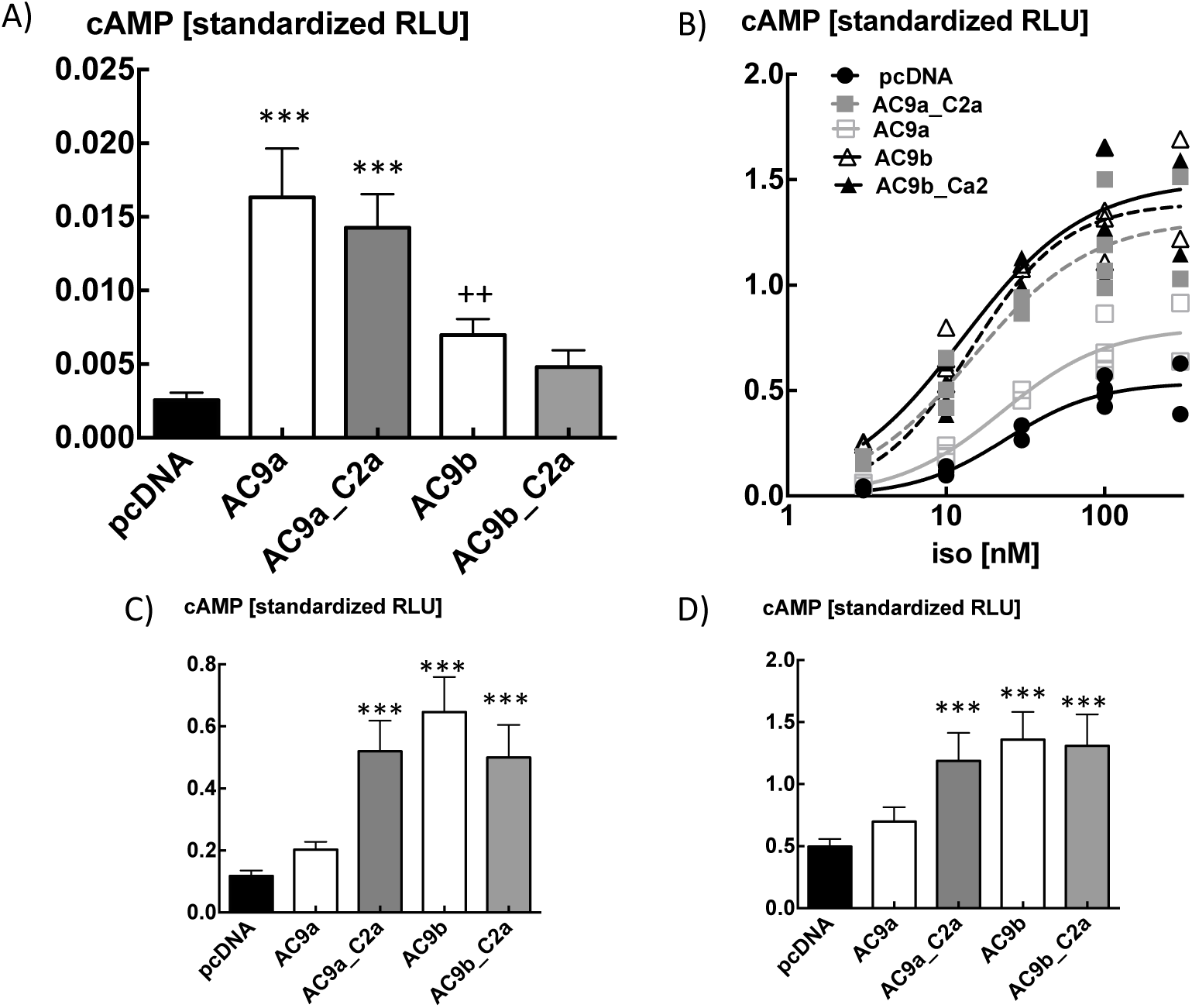
Distinct properties of zebrafish AC9a and AC9b, heterologously expressed in HEK293FT cells. AC9a_C2a and AC9b_C2a lack the C2b domain of the respective AC9a and AC9b proteins. A) Intracellular basal levels of cAMP, means±S.D., n=12 /group representative of three independent experiments with similar results. *** P<0.001 different from respective group of AC9b, ++P<0.01, different from AC9b_C2a. One-way ANOVA followed by Tukey’s multiple comparison test. B) Isoproterenol evoked intracellular cAMP levels, all data points are shown. Representative of three independent experiments with similar results. C) The cAMP response to 10 and D) 100 nM isoproterenol from B) — means±S.D., n=4/group. *** P<0.001 different from AC9. One-way ANOVA followed by Tukey’s multiple comparison test.

A yet further aspect of the ARM in human AC9 is autostimulation (Chen and Antoni 2023). This is likely due to the interaction of the first six N-terminal amino acid residues of ARM with the forskolin binding-pocket of AC9 (Qi et al. 2019; Nomura et al. 2025). However, the removal of C2b had only a relatively minor effect on the high basal levels of cAMP produced by AC9a while the reduction for AC9b was more substantial (basal cAMP levels generated by AC9a_C2a were 80±9 % of those generated by AC9a, n=3 independent experiments; for AC9b vs AC9b_C2a, this value was 45±9 %, n=4 independent experiments, data are means ±S.D., P<0.01 Student’s t-test). In the case of AC9b the greater reduction could be due to its relatively low basal activity in HEK293FTcells that is close to the instrument background.

## DISCUSSION

We found that a long C2b domain in *adcy9* is uniformly present and specific to vertebrate species. Gene duplication at TGD resulted in functionally distinct *adcy9* ohnologues that persisted in the teleost genome. The key trait that differentiates the sub-functionalized AC9 genes is autoinhibition of receptor stimulated cAMP production by the C2b domain. The respective expression patterns of the two AC9 mRNAs in post-TGD species indicate that autoinhibition is fundamentally important for brain function.

The amino-acid compositions of the C2b domains of the human as well as the two zebrafish AC9 isoforms clearly suggest intrinsic disorder (suppl. Fig. 1). This conforms to the cryo-EM studies of the bovine and human orthologues that failed to assign any structural features to the C2b domain of the protein (Qi et al. 2019; Nomura et al. 2025). Intrinsically disordered C-termini in proteins are frequently associated with autoinhibition (Trudeau et al. 2013; Uversky 2013). Indeed, a highly conserved and functionally relevant autoregulatory motif is invariably found in the C2b domain of vertebrate AC9a orthologs and appears to exert a profound autoinhibitory effect on receptor stimulated cAMP production (Pálvölgyi et al. 2018; Qi et al. 2019; Chen and Antoni 2023). There are various potential mechanisms of genome modification by which the C-terminal extension of *adcy9*, a gene that is already discernible in *C. elegans* (Antoni 2020), may have occurred (Herrera-Ubeda and Garcia-Fernandez 2021). Alternative exon splicing, recruitment of non-coding DNA seem unlikely as no significant similarity in the existing databases could be found out-with the *adcy9* genes of vertebrates. Transposon mediated transfer (Feschotte and Pritham 2007) or viral sequence domestication (Luebbert et al. 2025) seem more plausible but require further investigation.

Importantly, the long C2b tail of *adcy9* is present in cyclostomes that are the most ancient vertebrates alive today and already have a regionally differentiated brain (Sugahara et al. 2016). Indeed, similarly to rodent and human brain (Antoni et al. 1998; Visel et al. 2006; Sanabra and Mengod 2011) *adcy9* mRNA is widely expressed in lamprey brain see https://lampreybrain.kaessmannlab.org/ (Lamanna et al. 2023). When compared to the expression in the brain, only low levels of *adcy9* mRNA are found in the peripheral tissues of post-TGD teleosts, including zebrafish. Intriguingly, the ohnologue adcy9b has no ARM and the expressed protein showed no autoinhibitory activity with respect to receptor-stimulated cAMP production. Based on mRNA expression results in the PhyloFish database, the *adcy9b* gene has largely replaced *adcy9a* in all tissues except the brain. Proteomic analysis reported the presence of AC9a in the synaptosomal fraction of adult zebrafish brain (Bayes et al. 2017). Taken together with the functional analysis detailed in the present study, the data point to the subfunctionalization of the *adcy9* ohnologues in post-TGD teleosts.

Overall, little is known about the biological role of the *adcy9* gene. In rodents, deletion of *adcy9* causes marked (≈90%) pre-weaning lethality of the offspring homozygous for the deletion (Li et al. 2017; Perez-Garcia et al. 2018). The adult survivors show relatively minor defects of heart function and normal levels of anxiety (Li et al. 2017; Baldwin et al. 2022). In zebrafish, morpholino-based knock-down of *adcy9b* produced 50% mortality of embryos at birth and severe heart failure at 3 d.p.f. (Wu et al. 2020). Provided the morpholino effect was selective (Stainier et al. 2017), the results support the findings derived from the PhyloFish database as well as from real-time PCR in the present study, that *adcy9b* expression predominates in the heart. In this context it is of interest to note that a lower molecular weight AC9 that lacks the C2b domain is the dominant AC9 protein in the rodent heart and is also readily detectable in human heart tissue samples (Pálvölgyi et al. 2018; Rautureau et al. 2023). Similar to zebrafish AC9b, the truncated form of human AC9 has low basal activity and responds robustly to activation by Gs coupled receptors (Pálvölgyi et al. 2018; Chen and Antoni 2023). This indicates that autoinhibition of AC9 is not advantageous for heart function in zebrafish or mammals. As autoinhibition increases the Km for ATP by 40-fold (Qi et al. 2019), it seems possible that cAMP production by AC9 may be impaired at low levels of ATP (Rhana et al. 2024) that occur during the heart cycle. In turn, this would be detrimental for the pumping action of the heart (Zaccolo and Kovanich 2025). A further indication that autoinhibition in the single mammalian *adcy9* gene is an evolutionary burden in peripheral tissues is that the ARM is subject to dynamic phosphorylation at Ser1269 as well as Ser1273 (Barez-Lopez et al. 2023; Jiang et al. 2025). Indeed, alanine substitution of Ser1273 attenuated the autoinhibitory effect in HEK293 cells (Pálvölgyi et al. 2018).

A marked autostimulatory activity of C2b is readily demonstrated in human AC9 (Chen and Antoni 2023). In contrast, removal of the C2b domain from zebrafish AC9a had only a modest effect on basal cAMP levels. A possible explanation of this is that the zebrafish enzyme was tested in a human cell line, and the level of autostimulatory activity may depend on a species-specific cell context. Furthermore, zebrafish AC9a has two notable amino-acid changes when compared to its mammalian as well as its pre-TGD orthologs. First, V1266A of the highly conserved ARM is likely to weaken the interaction of ARM with the forskolin binding-pocket that is a positive allosteric modulator of AC9 (Qi et al. 2019; Qi et al. 2022). Second, W1160G is located within a short AC9-specific motif of the catalytic C2a domain that interacts with Gsα in the bovine enzyme (Qi et al. 2019; Qi et al. 2022). Thus, it seems possible that these two mutations in zebrafish AC9a, which are shared by the vast majority of the post-TGD AC9a orthologues currently in the databases, diminished the autostimulatory activity of C2b. The AC9b of zebrafish consistently produced lower basal cAMP levels than AC9a. As these were at best three-fold above the basal cAMP levels generated by the endogenous ACs of the host cell, the 50% reduction of basal levels observed upon removing the C2b domain translates into a marked, 75-80 % reduction of basal cAMP production when taking into account the output of the host cells. It is tempting to suggest that this is as yet a further aspect of subfunctionalization of the *adcy9* genes i.e. the segregation of autostimulation to *adcy9b* and autoinhibition to *adcy9a*.

In summary, TGD resulted in sub-functionalized *adcy9* ohnologs in teleosts to overcome an adaptive challenge met by post-translational modifications of the AC9 protein in mammals (Antoni 2024). On average, over 80% of ohnologue pairs revert to a single gene after TGD, thus the persistence of the *adcy9* ohnologs for over 200 million years indicates a significant adaptive advantage (Furutani-Seiki and Wittbrodt 2004; Braasch and Postlethwait 2012; Pasquier et al. 2017). The high expression of *adcy9a* in the brain and its restricted expression in peripheral tissues argues that the autoinhibitory feature is essential for some aspect of brain function or development. However, autoinhibition appears to be an evolutionary burden in peripheral tissues, particularly the heart. Hence, our results also illustrate the power of whole genome duplication to eliminate evolutionary burden.

## Data availability

All raw data and results are available from FAA upon request. The coding sequence of zebrafish *adcy9a* has been registered at Genbank as adenylyl_cyclase_9a accession no. PX511566.

## Acknowledgments

We thank Dr. Mark O. Collins, University of Sheffield for the reanalysis of proteomic data (Bayes et al. 2017).

## Author contributions

F.A.A. concept, design, data acquisition and analysis, wrote the paper, J. M. design, data acquisition and analysis, wrote the paper, H.M. design, data acquisition and analysis, wrote the paper, Z. C. design, data acquisition and analysis, L.Sz. design, data acquisition and analysis,, C. X. design, data acquisition and analysis, S. I. design, data acquisition and analysis, M. D.^,^ concept, design, data analysis, wrote the paper, A. B. concept, design, data analysis, wrote the paper, D.S. concept, design, data analysis, wrote the paper, M. J. S. concept, data analysis, wrote the paper, P.S. design, data acquisition and analysis, wrote the paper.

